# Single-molecule visualization of the effects of ionic strength and crowding on structure-mediated interactions in supercoiled DNA molecules

**DOI:** 10.1101/591008

**Authors:** Shane Scott, Cynthia Shaheen, Brendon McGuinness, Kimberly Metera, Fedor Kouzine, David Levens, Craig J. Benham, Sabrina Leslie

**Affiliations:** Department of Physics, McGill University, Montreal, Quebec, Canada H3A 2T8; Center for Cancer Research, National Cancer Institute, Bethesda, Maryland 20892; Genome Center, University of California Davis, Davis, California 95616

## Abstract

DNA unwinding is an important cellular process involved in DNA replication, transcription and repair. In cells, molecular crowding caused by the presence of organelles, proteins, and other molecules affects numerous internal cellular structures. Here, we visualize plasmid DNA unwinding and binding dynamics to an oligonucleotide probe as functions of ionic strength, crowding agent concentration, and crowding agent species using single-molecule CLiC microscopy. We demonstrate increased probe-plasmid interaction over time with increasing concentration of 8 kDa polyethylene glycol (PEG), a crowding agent. We show decreased probe-plasmid interactions as ionic strength is increased without crowding. However, when crowding is introduced via 10% 8 kDa PEG, interactions between plasmids and oligos are enhanced. This is beyond what is expected for normal in vitro conditions, and may be a critically important, but as of yet unknown, factor in DNA’s proper biological function in vivo. Our results show that crowding has a strong effect on the initial concentration of unwound plasmids. In the dilute conditions used in these experiments, crowding does not impact probe-plasmid interactions once the site is unwound.

## INTRODUCTION

Biological sciences have long sought to mimic in vivo conditions in vitro. Emulating cellular environments with current techniques remains a significant challenge due to the complexity of living cells. Cellular DNA, for example, is exposed to a wide variety of solutes of various sizes and physical properties including dissolved salts, proteins, and lipids (1). On average, 30% of the cell is occupied by solute molecules, though locally this value can rise to as high as 40% (1, 2). Current thinking in biology is that cells are highly compartmentalized, concentrating specific groups of molecules into droplet-like formations, enhancing this crowding effect (3). This so-called crowding may have an effect on cellular interactions and structures (4, 5).

Additionally, the cytoplasm and its contents are confined by the cellular membrane to a small space of a few micrometers for bacteria, and around one hundred micrometers in eukaryotic cells. DNA and other nuclear materials are themselves confined within the eukaryotic nuclear membrane, an even smaller confined region than the cytoplasm. Both the crowding created by the molecules and the confinement afforded by the membrane are hard to emulate and are thus neglected in standard in vitro studies. However, molecular crowding, where dissolved substances may act as obstacles to reactions and cellular structures, and confinement, where the cellular machinery is forced into small spaces, may impact biological function and are of strong interest to the scientific community (4, 6, 7).

A number of cellular structures - from organelles, to cellular scaffolding, to DNA molecules - are affected by crowding, ionic concentration, and confinement. Small, dissolved ions can screen repelling charges on molecules, such as the negatively-charged phosphates along the DNA backbone. By influencing molecular hydration, solution viscosity, and intermolecular collisions, crowding can affect cellular structures and macromolecules, such as enzymes and other proteins (4). Collisions between crowding agents and reactants can either inhibit or enhance reactions, depending on the system under consideration (5, 8). On the one hand, molecular diffusion is reduced due to collisions with crowders, which can impede reactants from colliding and reacting. On the other hand, crowding agents can force reactants to encounter one another more often by ‘caging’ them together, leading to an increase in reaction rates (5, 9). To investigate these effects, polyethylene glycol (PEG), dextran, polyvinylpyrrolidone (PVP), and molecules found in cells, such as DNA, have all been used to imitate crowding conditions in vitro (1, 10).

Environmental conditions within cells are particularly important when considering higher-order DNA structure created by supercoiling, the torsional stress applied to DNA. This torsional tension, caused by under- or over-twisting the DNA double helix, has been shown to drive structural transitions in DNA, such as those from B-form to Z-form, or to fully unwound (11, 12, 13). These transitions are important physiologically: once the DNA is in one of these alternate structures, cells may initiate important processes which require unwinding of the double-helix, such as DNA transcription and replication (14, 15). Understanding the role the solution environment plays on supercoiled DNA is thus important for understanding cellular processes. For example, supercoiling decreases with increasing salt concentration: salt reduces electrostatic repulsion, decreasing the persistence length and thus the supercoiling free energy (16, 17). DNA secondary structure, such as site-unwinding, is affected by changes in ionic strength: a reduction in electrostatic repulsion and supercoil-derived free energy can reduce the probability of unwinding (12).

The presence, or absence, of solute molecules as steric obstacles is also important to DNA structure formation (2, 6). Crowding has been shown to have affect the unwinding of two oligonucleotide strands (18, 19), on hairpin formation in single-stranded DNA (ssDNA) oligos (2, 18), and on G-quadruplex formation (20). Recent work has also shown the importance of crowding and confinement on DNA structure (21, 22). Molecular dynamics simulations have demonstrated decreasing DNA duplex stability with increasing crowding agent size and concentration (23). How both crowding and ionic strength impact DNA secondary structure in more complex systems, such as supercoiled plasmids, is still unclear.

The local unwinding of DNA is critical to proper cellular function: in cells, many DNA-DNA and DNA-protein interactions require the double-stranded DNA (dsDNA) to be unwound before they can occur. Site-specific unwinding can thus regulate gene expression by allowing proteins access to these locations along a DNA molecule (24). Homologous recombination is also nucleated by this unwinding, with the opening of the dsDNA allowing single-stranded DNA to bind (25, 26). Gene editing techniques, such as CRISPR-Cas9, involve interactions between an RNA oligo and a target DNA site. DNA unwinding is in turn controlled in part by the presence of salts and crowding agents. A more complete understanding of supercoil-induced unwinding would thus provide important insights into these processes, detailing mechanisms by which the cellular environment, including local ionic strength and crowding, will affect processes occurring within cells.

We recently reported a novel approach to study DNA unwinding in which the binding of a fluorescently-labeled oligonucleotide probe to the plasmid’s known unwinding site was used to measure its open state (13). By measuring the diffusion coefficient of the fluorescently-tagged probes, it was possible to determine when they were bound to plasmid unwinding sites. Sample molecules were confined to an array of glass nanopits using the CLiC technique and imaged at multiple points over the course of two hours. Pit arrays in the flow cells allowed for multiple, independent reaction chambers to be monitored over long time periods. The pits we used are on the same size scale (3 *µ*m diameter, 500 nm deep) as an E. coli cell (500 nm diameter, 3 *µ*m long). In this previous study, we demonstrated how unwinding and reaction rate between the probe and plasmid increased with temperature, but only unwinding was affected by supercoiling (13).

Here, we build off of the previous work by exploring DNA unwinding using a range of in vitro conditions that approach those found in cells. Specifically, we introduce crowding agents, mimicking the cell’s crowded environment, and vary the ionic strength to observe these important effects on reaction rates and DNA unwinding.

## MATERIALS AND METHODS

### Oligo probes

A preliminary prediction of the Site 1 unwinding region on the pUC19 plasmid at our experimental conditions was made with the computational algorithm given by Zhabinskaya and Benham (12). Applying these results, we designed a DNA probe oligo to be complementary to the known unwinding region of pUC19 with the sequence 5’-GAT TAT CAA AAA GGA TCT TCA CCT AGA TCC-3’, and ordered it from IDT (San Jose, CA, USA), including an amino C6 linker group on the 5’ end. To visualize its probe binding to the pUC19 unwinding site, we use this amino linker to fluorescently tag the probes with a Cy3B NHS ester purchased from GE Healthcare (Mississauga, ON, Canada) and dissolved at a concentration of 1 mg/mL in dimethyl sulfoxide (DMSO). As the diffusion of the free probes is much higher compared to that of the pUC19 plasmids, we use a diffusion-based assay to determine whether or not the probes are interacting with the plasmid’s unwinding site. By visualizing the fluorescence and tracking the intensity peak over time, we can differentiate probes bound to slow-moving plasmids from the quickly diffusing free probes.

Dried probes were dissolved at 20 mg/mL in 0.1 M NaHCO_3_ with pH 8.4, and mixed overnight with an excess of Cy3B dye. To remove unreacted dye, ammonium acetate was added to the probes followed by 3 washes with ethanol to precipitate it. Dried probes were dissolved in 10 mM Tris (pH 8.0), and Cy3B-labeled DNA was purified from unlabeled probes using a C18 reverse phase column on a high performance liquid chromatography (HPLC) device.

### Plasmid topoisomers

DH5*α E. coli* cells were transformed with pUC19 from New England Biolabs (Whitby, ON, Canada). Cells were grown in 1 × LB media overnight, and plasmids were harvested from this growth using a Qiafilter Plasmid Midi Kit from Qiagen Inc. - Canada (Toronto, ON, Canada). Plasmids were supercoiled using the protocol by Keller by reacting pUC19 with topoisomerase IB from Thermo Fisher Scientific (Toronto, ON, Canada) in the presence of ethidium bromide (27). Purification of supercoiled plasmids was performed using a Qiagen PCR Purification Kit from Qiagen Inc. - Canada (Toronto, ON, Canada).

The superhelical density of pUC19 topoisomers was verified using a 2% agarose gel with 10 mM tris (pH 8.0), 20 mM sodium acetate, 1 mM ethylenediaminetetraacetic acid (EDTA) and either 3 mg/mL or 6 mg/mL chloroquine diphosphate run at 30 V for 40 h. Gels were stained with SYBR Safe from Thermo Fisher Scientific and imaged with a Bio-Rad (Mississauga, ON) GelDoc EZ Imaging System. Topoisomer samples demonstrated a Gaussian distribution of superhelical densities, which was analyzed using Matlab software to obtain the mean superhelical density ⟨*σ*⟩ and standard deviation *s*_*σ*_.

Three distributions of topoisomers centered around ⟨*σ*⟩ = −0.07, −0.101, and −0.132 were used throughout this work. Supercoiling increases as the absolute value for superhelical density increases away from 0, so a sample with ⟨*σ*⟩ = −0.132 is more supercoiled than one with ⟨*σ*⟩ = −0.07. When extracted from cells, topoisomers have a distribution of supercoiling values centered around ⟨*σ*⟩ = −0.055, though cells can further supercoil their DNA to promote unwinding (28). As shown in Scott, *et al.*, for the system presented in this work, increasing the overall supercoiling promotes binding due to an increase in site-unwinding (13).

### Convex lens-induced confinement (CLiC) microscopy

The microscopy setup, data acquisition, and image analysis were all performed using CLiC as in Scott, *et al.* (13). In this microscopy system, two coverslips are separated by 30 *µ*m-thick double-sided tape. A lens is used to compress the top coverslip, forcing it into contact with the bottom coverslip, which possesses an array of 3 *µ*m-diameter by 500 nm-deep pits. Compressing the coverslips allows molecules to be held within the field of view while simultaneously allowing them to diffuse freely within the pits. Sample videos were taken every minute over ~2 hours with 50 ms exposure time at 37°C. Binding events were identified using an image-processing algorithm as described in Scott, *et al.* (13). All samples were dissolved in water with 12 mM Tris, 25 mM HEPES, 2.5 mM protocatechuic acid (PCA), and 50 nM protocatechuate 3,4-dioxygenase (PCD) at a pH of 8.0. The combination of PCA and PCD act as an oxygen scavenging system that impedes fluorophore photobleaching.

To estimate binding rates and determine the unwound state of plasmids, data sets were fit to a second-order reaction model that assumes the rates of unwinding and rewinding are negligible compared to the rate of binding, consistent with prior results (13). As the probe molecules are in excess to the bound complexes for all experimental conditions, the second-order reaction equation can be expressed as

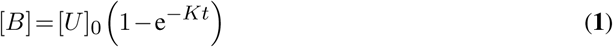

where *K* is a constant, *t* is the time, [*U*]_0_ is the initial concentration of unwound plasmids, and [*B*] is the concentration of bound complexes. The constant *K* can be used to determine the rate of reaction through *K* = *k* ([*P*]_0_ − [*U*]_0_), where *k* is the true rate constant for this reaction, and [*P*]_0_ is the initial concentration of probe molecules (13).

## RESULTS

### Crowding Agents Affect DNA Binding and Unwinding

Crowding agents are assumed to influence binding events mostly through collisions with other molecules. It is important to assess whether crowding agents exert specific effects on binding, especially at physiological ionic strengths. Molecules that interfere in this manner are unsuitable for use as crowding agents. To investigate these effects, experiments with three crowding agents (8 kDa PEG, 10 kDa PVP, and 55 kDa PVP) were performed at 12.5% (w/v) with an ionic concentration of 150 mM (137.5 mM NaCl, 12 mM Tris, 25 mM HEPES, pH 8.0) and temperature of 37°C (Fig. 1). At pH 8.0, only NaCl ions dissociate completely, leading to an ionic contribution of 12.5 mM from both Tris and HEPES. An increase in binding is observed for all experiments with crowding agents compared to a control with no crowding agent. The increase in binding is similar between the specific species and sizes of crowding agents tested. As there wasn’t a significant difference in binding caused by a different species or size of crowding agent, it is probable that none of the crowding agents tested affect the plasmid-probe system either electrostatically or through hydrophobic interactions. In subsequent experiments, we used 8 kDa PEG as the crowding agent.

**Figure 1.**
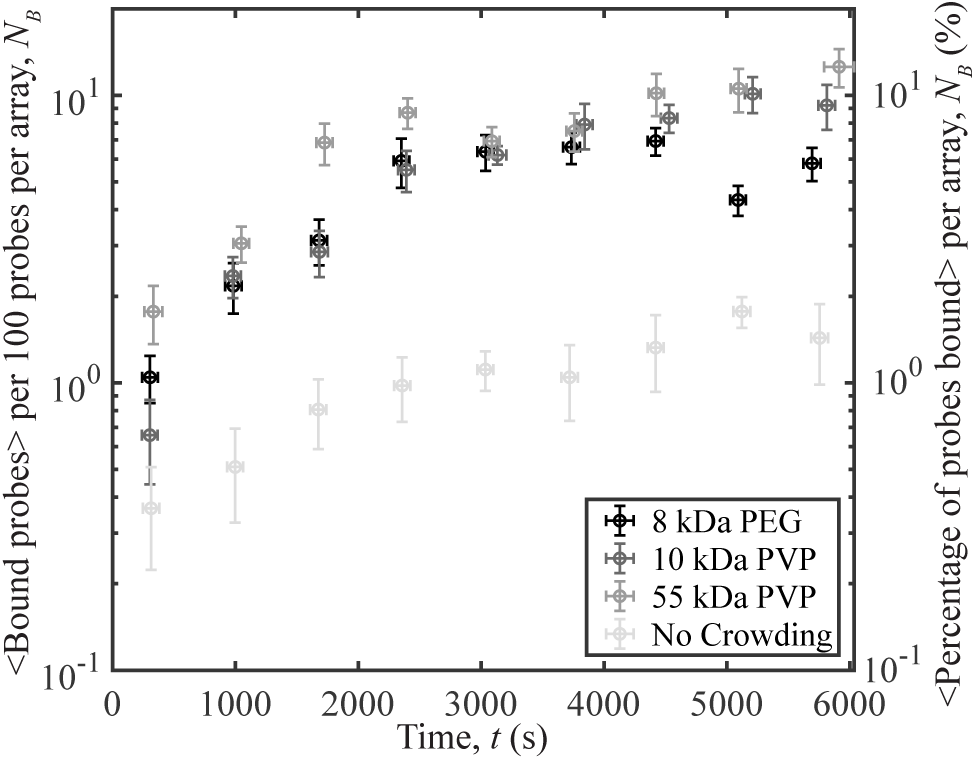
Probe-plasmid binding and different crowding agents at 12.5% (w/v). The number of binding events per 100 probes averaged over 10 different videos vs. time, with a selection of crowding agents. The plasmids have a superhelical density of ⟨*σ*⟩ = −0.132, *s*_*σ*_ = 0.007. To distinguish between the unbound probes, and probes bound to plasmids, experiments and image analysis were performed as described in the Supplementary Information. Error bars denote the standard error of the mean of the number of bound probes on the y-axis, while those on the x-axis represent the standard error of the mean of the measurement times.

### Binding Increases With Crowding Agent Concentration

To investigate how crowding affects DNA-DNA interactions and plasmid unwinding, pUC19 with ⟨*σ*⟩ = −0.101 and *s*_*σ*_ = 0.004 were reacted with probe molecules in the presence of increasing concentrations of 8 kDa PEG, 150 mM ionic strength, and temperature of 37°C. The range of crowding agents used in these experiments, 0%, 1%, 5%, 10%, and 20% (w/v), correspond to excluded volumes of 0%, 1.31%, 6.55%, 13.1%, and 26.2% (v/v), respectively. See the Supplementary Information for calculations of the excluded volume. We observe an increase in binding with increasing 8 kDa PEG concentration as shown in Fig. 2 a). This is primarily due to an increase in unwound sites according to the fits obtained from our model, as in Fig. 2 b). Note that at a PEG concentration of 20%, the crowding agent concentration was sufficiently high to reduce the diffusion of the probes so that they could be tracked using a 50-ms exposure time. To distinguish between the probes and plasmids, we determined a new threshold diffusion coefficient value, where all molecules diffusing below this were considered bound probe-plasmid complexes. Details for this procedure are found in Supplementary Information.

**Figure 2.**
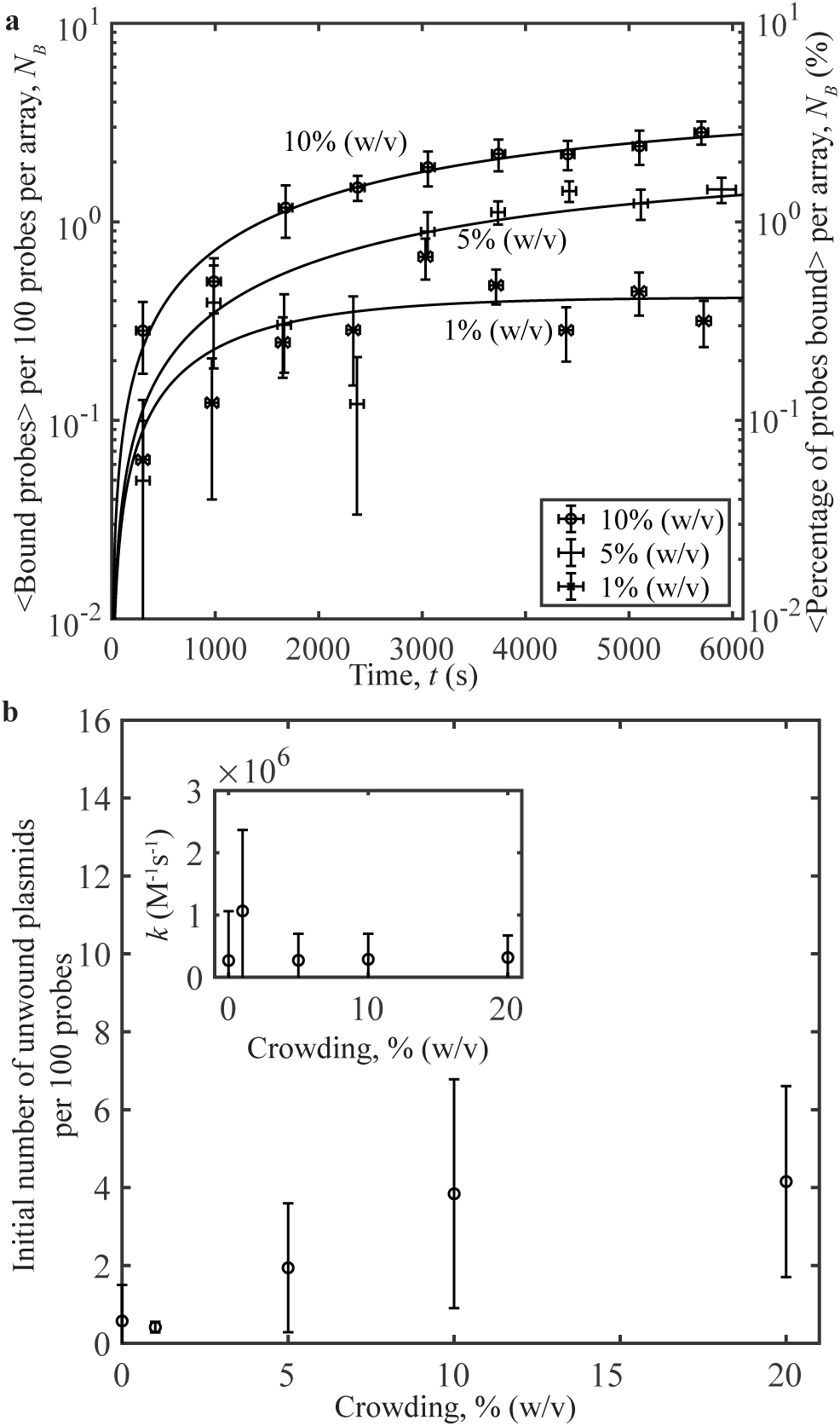
Probe-plasmid binding and crowding agent concentration. **a)** The number of binding events per 100 probes averaged over 10 different videos vs. time, for three concentrations of 8 kDa PEG. Curves represent fits to the data. The plasmids have a superhelical density of ⟨*σ*⟩ = −0.101, *s*_*σ*_ = 0.004. **b)** Initial amount of unwound plasmid obtained from the fits in a) per 100 probes vs. concentration (w/v). Inset: Binding rates to unwound sites vs. concentration obtained from the data fits. Error bars were calculated as in Fig. 1.

### Microscopy of Salt-dependent Interactions

Unwinding predictions vs. salt concentration were made for Site 1 on a pUC19 plasmid with ⟨*σ*⟩ = −0.101 and *s*_*σ*_ = 0.004 using the DZCBtrans algorithm created by Zhabinskaya and Benham (Fig. 3) (12). As ionic concentration is increased, the number of unwound bases in Site 1 decreases rapidly.

**Figure 3.**
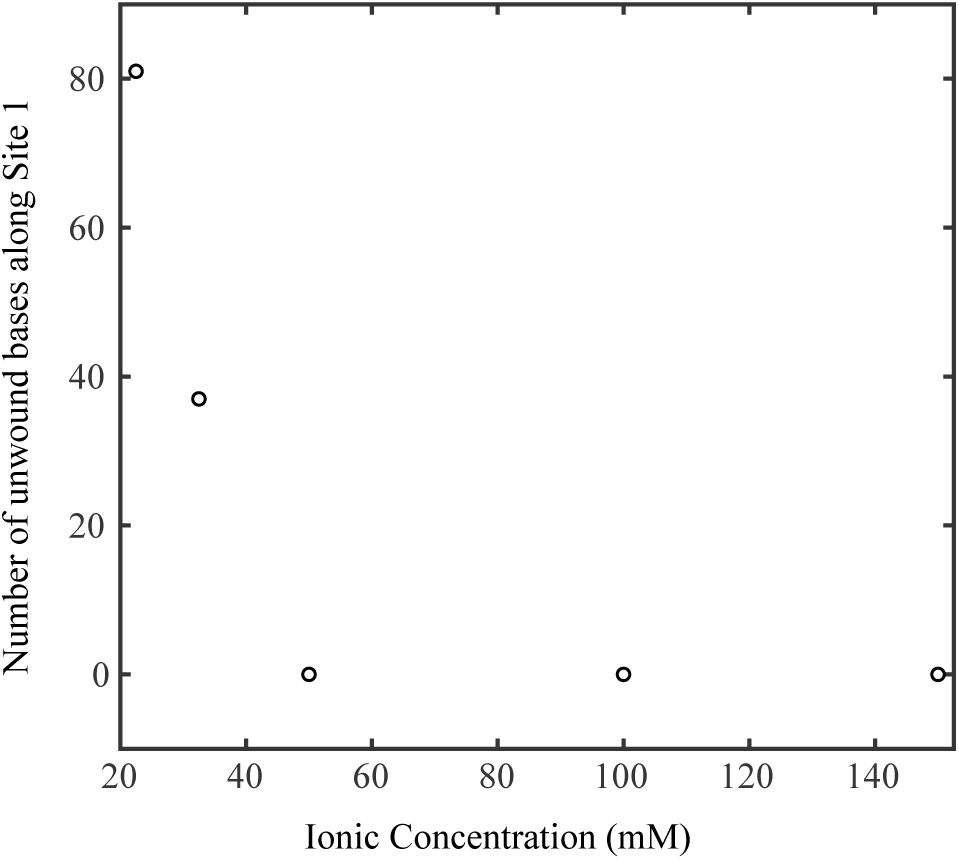
Predicted number of unwound bases along Site 1 vs. ionic strength. Numerical simulations were performed for pUC19 with ⟨*σ*⟩ = −0.101, with *s*_*σ*_ = 0.004 (12). As ionic strength increases, the number of bases along Site 1 is predicted to decrease.

A range of 10 mM ≤ [NaCl] ≤ 137.5 mM was used in experiments with pUC19 at a constant ⟨*σ*⟩ = −0.101 and *s*_*σ*_ = 0.004 to observe the effects of monovalent ionic concentration on plasmid-probe binding (Fig. 4a). Taking into account the ionic contributions of NaCl with those of the 12 mM Tris and 25 mM HEPES present (≈12.5 mM total contribution from dissociated ions at a pH of 8.0) provided a total ionic concentration range of 22.5 mM - 150 mM. Concentrations between 100-200 mM are considered to be physiological ionic conditions (29). As salt concentration is increased, a decrease in binding occurs. We expect binding to decrease with increasing ionic strength because an increase in positively charged ions screens the negative charges on the DNA. This renders unwinding in pUC19, and thus probe binding, to be less likely. Increased ionic strength also decreases the supercoiling energy (17), thereby reducing the probability of unwinding, which would also lead to a decrease in probe binding.

**Figure 4.**
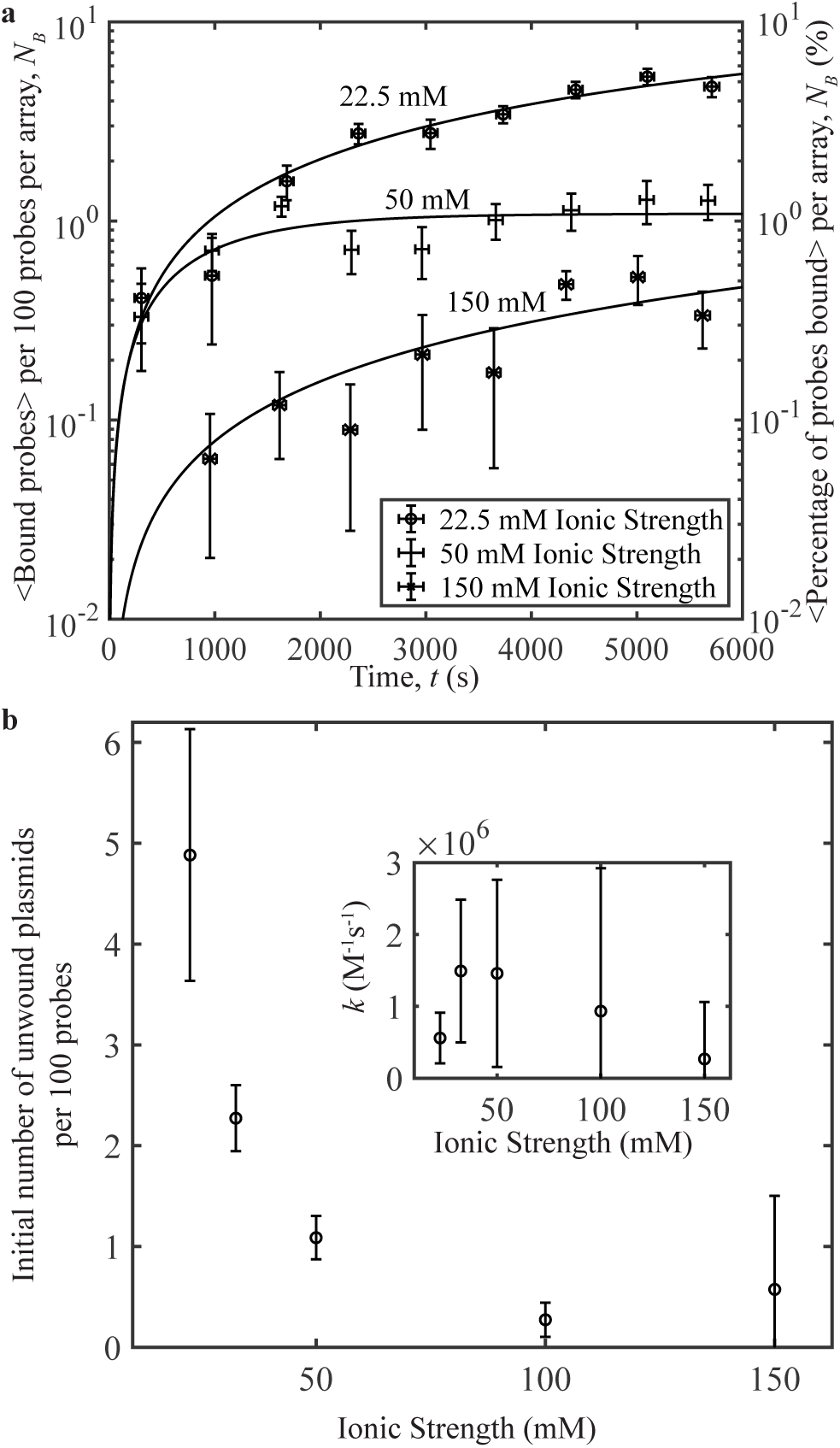
Probe-plasmid binding and ionic strength. **a)** The number of binding events per 100 probes averaged over 10 different videos vs. time, for three ionic strengths studied. Curves represent fits to the data. The plasmids have a superhelical density of ⟨*σ*⟩ = −0.101, *s*_*σ*_ = 0.004. **b)** Initial number of unwound plasmid sites obtained from the fits in a) per 100 probes vs. ionic strength. Inset: Binding rates to unwound sites vs. ionic strength obtained from the data fits. Each data point was averaged over ~3,300 probes. Error bars were calculated as in Fig. 1.

To determine what exactly accounted for the reduction in binding, whether it be a decrease in unwinding, a reduction in the reaction rate between the plasmid and probe, or both, we plotted the initial amount of unwound plasmid and the rate of binding vs. ionic strength. These values were obtained by fitting the data to Eq. **1** and extracting the fit parameters from the line of best fit. From the fits, the initial amount of unwound plasmid decreases with increasing ionic strength, while the rate of binding to unwound sites does not vary significantly for all ionic strengths observed when the errors on the binding rates are considered (Fig. 4 b). This agrees with the unwinding predictions presented in Fig. 3.

### Binding Increases with Ionic Concentration in the Presence of Crowding Agents

To test the effects of ionic strength in the presence of crowding agents, pUC19 samples with ⟨*σ*⟩ = −0.101, *s*_*σ*_ = 0.004 were reacted with probes in the presence of 10% (w/v) 8 kDa PEG at 37°C and increasing NaCl concentration. As in the previous section, a range of 10 mM ≤ [NaCl] ≤ 137.5 mM was used, providing a total ionic strength range of 22.5 mM - 150 mM when the ionic contributions of the 12 mM Tris and 25 mM HEPES were added at pH 8.0. In the presence of crowding, binding between the probe and plasmid increased with increasing salt concentration (Fig. 5), in contrast to the results presented in Fig. 4. The fits to Eq. **1** presented in Fig. 5 b) compare the initial concentrations of unwound pUC19 and reaction rates vs. ionic strength for the samples reacted with and without 10% (w/v) 8 kDa PEG. The reaction rate is unaffected by the presence or absence of crowding agents. Additionally, unwinding decreases as ionic strength increases when no 8 kDa PEG is in the system, but increases with ionic strength when the crowding agent is present.

**Figure 5.**
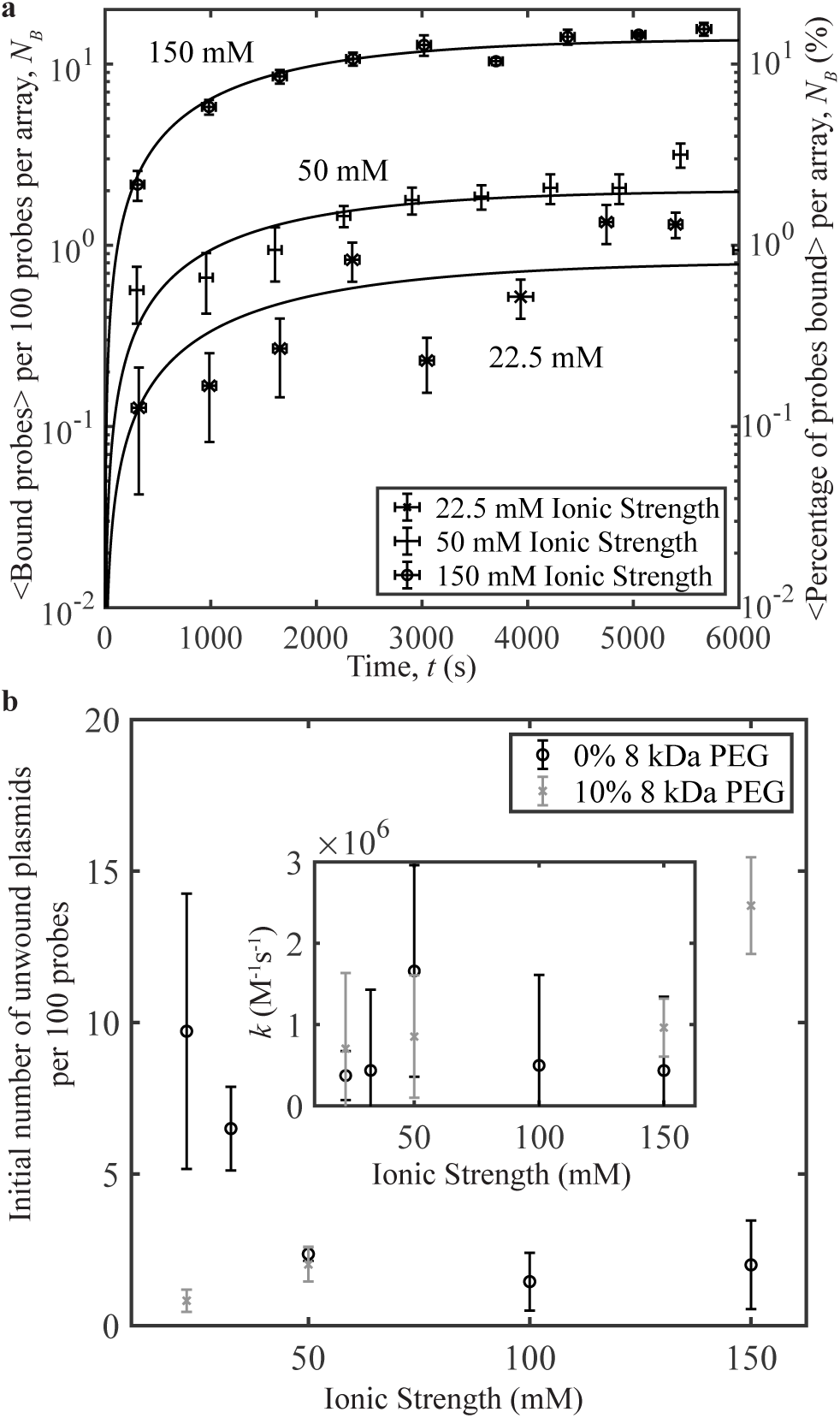
Probe-plasmid binding and ionic strength with and without 10% 8 kDa PEG at. ⟨*σ*⟩ = −0.101**. a)** The number of binding events per 100 probes averaged over 10 different videos vs. time, for three ionic strength concentrations. Curves represent fits to the data. The plasmids have a superhelical density of ⟨*σ*⟩ = −0.101, *s*_*σ*_ = 0.004. **b)** Initial number of unwound sites obtained from the fits in a) and Fig. 4 a) per 100 probes vs. ionic strength. Inset: Reaction rates vs. ionic strength obtained from the fits in and Fig. 4 a). Each data point was averaged over ~3,300 probes. Error bars were calculated as in Fig. 1.

To verify that the increase in binding with ionic strength in the presence of crowding can occur for a range of supercoiling, we repeated the experiment above using pUC19 plasmids with a different superhelical density of ⟨*σ*⟩ = −0.07, *s*_*σ*_ = 0.007. The ionic range studied was exactly the same as in the previous experiments, with a total ionic concentration range of 22.5 mM - 150 mM. As with the previous sample that had higher supercoiling, as ionic strength increased, the amount of binding increased over time in the presence of crowding agents (Fig. 6). Plotting the parameters obtained from fits to Eq. **1** in Fig. 6 b), the initial concentration of unwound plasmids increases, indicating that the increase in binding is caused by an increase in the initial amount of unwound plasmid with increasing ionic strength. The subsequent rate of binding to the “open site” - once unwound - did not depend on ionic strength. The results from Fig. 6 confirm the trend seen in Fig. 5, and indicates that the results are unaffected by the difference in superhelical density.

**Figure 6.**
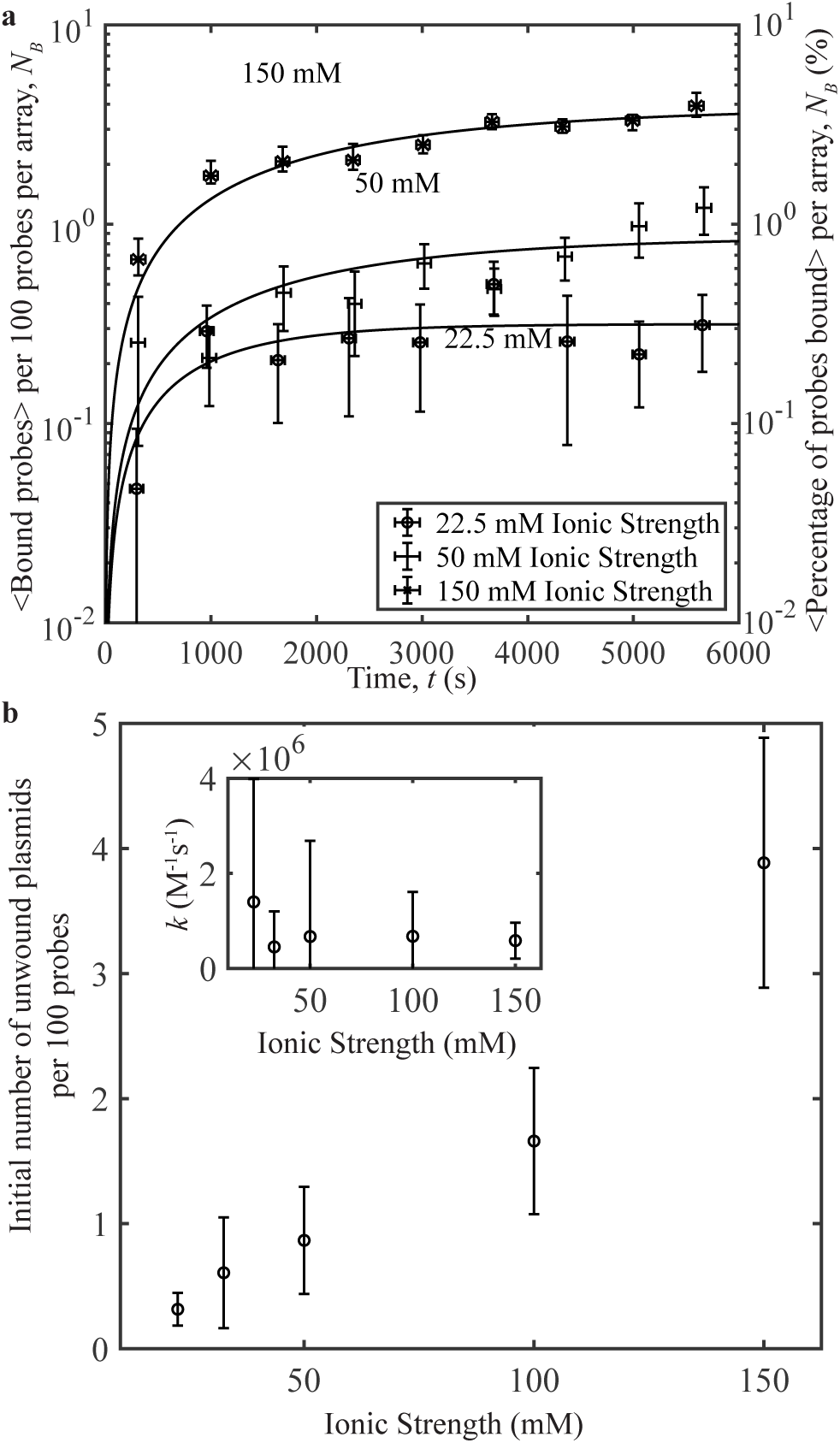
Probe-plasmid binding and ionic strength in the presence of 10% 8 kDa PEG at. ⟨*σ*⟩ = −0.07**. a)** The number of binding events per 100 probes averaged over 10 different videos vs. time, for three ionic strength concentrations. Curves represent fits to the data. The plasmids have a superhelical density of ⟨*σ*⟩ = −0.07, *s*_*σ*_ = 0.007. **b)** Initial number of unwound sites obtained from the fits in a) per 100 probes vs. ionic strength. Inset: Reaction rates vs. ionic strength obtained from the fits. Each data point was averaged over ~3,300 probes. Error bars were calculated as in Fig. 1.

At levels of crowding around 10% (w/v), DNA undergoes a phase transition to a tight, globular molecule at higher salt concentrations (30). While this compacts the supercoiled molecule further, it may liberate supercoiling energy normally used in writhe to overcome electrostatic repulsion from the negatively-charged phosphate backbone. This supercoiling energy would create further tension along the DNA molecule, which is energetically unfavored. To relieve this tension, it may be possible that the excess supercoiling energy is used for secondary structure formation; in the case of this system, DNA unwinding in pUC19. A more in-depth discussion of what may be occurring here is presented in the following section.

## DISCUSSION

The experimental and theoretical results presented in this work demonstrate surprisingly important effects of near-physiological salt and crowding conditions on supercoil-induced unwinding. When ionic strength is increased without crowding agents present, there is a marked decrease in probe-plasmid binding. Remarkably, when 10% 8 kDa PEG is introduced and salt concentrations are increased, we find that probe-plasmid binding increases, the complete opposite effect to the case without crowding. Importantly, this is primarily due to an increased number of unwound sites. This has implications in the study of all DNA-dependent processes in the crowded cellular and nuclear environments, such as in the cases of DNA repair and replication, whose dynamics rely on the conditions of the surrounding cellular environment.

In bacteria, whose physiological ionic conditions are within the 100-150 mM range and which can supercoil plasmids to ⟨*σ*⟩= −0.101, our data indicates that unwinding in pUC19 should be an unlikely event without crowding agents present. These conditions would prevent pUC19 replication as the unwinding site studied in this work is associated with the plasmid’s origin of replication, a strange effect considering that pUC19 is known to have a high copy number in *Escherichia coli* bacteria. We discover here that this low unwinding at physiological salt conditions may be counteracted by the presence of crowding agents, as shown in Fig. 2, 5, and 6. Eukaryotic cells also possess numerous non-B-DNA secondary structures, including unwound regions, as shown using potassium permanganate fingerprinting (31). These regions have been shown to have some regulatory potential, making it all the more important that the mechanisms of unwinding at physiological conditions be fully understood.

It is possible that the combined effects of crowding and elimination of electrostatic repulsion along the negatively-charged phosphate backbone could liberate further supercoiling energy, allowing for more unwinding. Previous work by Karimata *et al.* on the effects of salt concentration on DNA destabilization demonstrated that DNA unwinding in the presence of 8 kDa PEG should decrease with increasing ionic strength (32). Destabilization of the DNA in that work, however, focused on destabilizing small hairpins formed by complementary, short oligos, in contrast to the system presented here where unwinding is supercoil-driven.At 10% 8 kDa PEG and high ionic strengths, work by Zinchenko, *et al*. has shown that DNA should begin to compact into a globular, condensed state (30). This crowding-induced condensation has been shown to force DNA to adopt rigid, toroidal structures (33), which have also been observed with plasmids using atomic force microscopy (34). It is thought that these toroids are formed via interactions between DNA filaments (35), creating what looks to be DNA loops superimposed on themselves (34, 36). Work by Zinchenko, *et al.*, has also shown that the melting temperature of DNA strands may increase in DNA compacted into a toroidal form, similar to the unwinding effect that we show here (37). In our system, this may occur as the crowding forces the DNA plasmids into rigid, toroidal structures, lowering the number of available structural conformations. This structural restriction decreases the entropy of the system and increases its free structural energy. The increased energy may then be used to allow previously inaccessible, higher-energy structural transitions, such as DNA unwinding, to form. Furthermore, unpaired regions could act as hinge regions, further aiding in compaction and liberating even more supercoiling energy.

The crowding concentrations presented in this work are in a similar range to those in cells. In the nucleus, the concentration of crowding is thought to be between 20% and 40%, though the actual value varies locally and is difficult to measure, especially in eukaryotes (38). As our results lie in the range of nuclear crowding concentrations, they may have an impact on how DNA behaves in eukaryotes. Organelles, proteins, and other cellular machinery could contribute to crowding in cells, promoting unwinding, where high ionic strengths alone would otherwise suppress unwinding. That crowding may make DNA structural transitions occur with higher probability than one would expect is of particular importance as it may be crucial for the DNA to function properly in living cells.

Superhelical density has a significant impact on probe-plasmid binding due to its important effect on unwinding: experiments from our previous work in Scott, *et al.* (13) demonstrated less than 2% of probes bound at 22.5 mM ionic strength when ⟨*σ*⟩= −0.07. Comparing the binding in this work with that from the previous work, it can be seen that the presence of 10% 8 kDa PEG increased binding by a factor of 2. As the reaction rates between the probes and plasmids remained constant for all crowding agent concentrations, the increase in binding in the presence of 8 kDa PEG was mainly due to an increase in unwinding. The crowding effect on DNA unwinding is therefore comparable to that of supercoiling. This is a highly relevant finding for future studies of other DNA structures such as G-quadruplexes or DNA cruciforms, where cellular crowding is thought to play a significant role on structure formation (2, 18, 20).

An increase in the number of bound complexes with increasing 8 kDa PEG concentration reinforces the idea that crowding plays a key role in governing DNA structure. At the majority of the concentrations of 8 kDa PEG used in this work, the solution is in a semidilute regime, exceeding the crossover concentration *c*\* = *N* ^−4/5^ ≈ 4% (w/v). It is at this concentration that the PEG molecules begin to overlap with one another, the beginnings of a mesh-like network where the molecules entangle one another (39). At these concentrations, crowding agent molecules may interact with pUC19 plasmids by inserting themselves into, or otherwise stabilizing, their unwinding regions, physically preventing the site from closing, and leaving them available for probe binding. In vivo, cellular machinery could mimic this effect, specifically through molecules small enough to “fit” in-between the unwound DNA strands. This has implications for targeted gene editing techniques, such as CRISPR-Cas9, which employs a guiding RNA oligo strand to target specific regions of DNA. Cellular crowding would be expected to aid in the unwinding of target DNA and allow for the binding of the guide RNA to its target, despite high physiological ionic strengths which could impede target DNA opening. A better understanding of how a crowded environment affects reactions could also benefit researchers studying other kinds of biomolecular reactions, such as ligation (40, 41).

## CONCLUSION

In summary, we have used single-molecule CLiC microscopy to visualize and investigate the interactions between a probe oligo and a supercoil-induced unwinding site on pUC19 plasmids as functions of ionic strength and crowding effects. By increasing salt concentration, we have demonstrated a decrease in probe-plasmid binding, that, in turn, indicates a decrease in DNA unwinding at physiological salt conditions. The presence of crowding agents, such as 8 kDa PEG, counteracts this effect, augmenting probe-plasmid binding due to an increase in DNA unwinding at physiological ionic conditions. Verification that 8 kDa PEG does not interact with our system via electrostatic or hydrophobic interactions was achieved by establishing similar binding in the presence of 10 kDa and 55 kDa PVP, which also act as crowders. Increasing concentrations of 8 kDa PEG are demonstrated to *enhance* probe-plasmid binding and unwinding. In all conditions studied, we show that varying ionic strength and crowding agent concentration has a negligible effect on the rate of binding to unwound sites in the probe-plasmid system, indicating that increased binding is driven principally by an *increase* in unwinding. These studies have important widespread implications in the field of molecular biology and the study of DNA structural dynamics. Furthermore, the results presented in this work are helpful in the development of biotechnologies such as CRISPR-Cas9 which rely on the interactions of oligos with target DNA at physiological conditions, and a broad range of enzymatic reactions.

## ACKNOWLEDGEMENTS

The authors thank the Canadian Foundation for Innovation, the National Science and Engineering Research Council of Canada Discovery program, The Fonds de recherche du Quebec Nature et technologies (FRQNT) Team Grant and Fellowship programs, the Cellular Dynamics of Macromolecular Complexes CREATE fellowship program, the Bionanomachines fellowship program, and McGill University for funding and resources. C.S. thanks CMC Microsystems for fellowship funding for fabrication. The authors thank Tom Edwardson and Hanadi Sleiman for oligo purification on an HPLC.

