## Supplementary Information for "Single-molecule visualization of the effects of ionic strength and crowding on structure-mediated interactions in supercoiled DNA molecules"

February 26, 2019

1. Department of Physics, McGill University, Montreal, Quebec, Canada H3A 2T8
2. Center for Cancer Research, National Cancer Institute, Bethesda, Maryland 20892
3. Genome Center, University of California Davis, Davis, California 95616

#### Supplementary Information: Methods

##### Excluded Volume Calculation for 8 kDa PEG

To calculate the excluded volume in our solutions of 8 kDa PEG, it is necessary to determine the excluded volume of a single 8 kDa PEG molecule. Here, we assume that 8 kDa PEG acts as a real chain in solution, where each polymer is modelled as $N_K$ rigid rods of size $b_K$ connected together. The volume $v \approx R_F^3$ of a single 8 kDa PEG molecule can be estimated using the Flory radius $R_F = b_K N_K^{3/5}$. From Choi, *et al.*, $b_K = 0.7$ nm for PEG in a good solution [1]. We can calculate $N_K = R_{max}/b_K = Nb/b_K$, where $R_{max}$ is the contour length of a single 8 kDa PEG molecule, $N$ is the number of ethylene glycol monomers, and $b$ is the size of a monomer of ethylene glycol. From Choi, *et al.*, the approximate size of a single ethylene glycol molecule is $b = 0.44$ nm, with a molecular weight of 44 g/mol [1]. Thus, $N_K = (8,000/44)(0.44\text{ nm})/(0.7\text{ nm}) = 114.3$, $R_F = (0.7\text{ nm})(114.3)^{3/5} = 12$ nm, and the excluded volume is $v \approx (12\text{ nm})^3 \approx 1,700\text{ nm}^3$.

Using the excluded volume of one 8 kDa PEG molecule, the percent of excluded volume in solution can be calculated. One gram of 8 kDa PEG must have an excluded volume of $v_g = N_A v/(8,000\text{ g/mol}) \approx 131$ mL/g, where $N_A$ is Avogadro's number. Concentrations of 8 kDa PEG used in this work are 1%, 5%, 10%, and 20%, all in weight/volume, or grams per 100 mL of solution. The excluded volume concentrations are then 1.31%, 6.55%, 13.1%, and 26.2% (v/v) for the 1%, 5%, 10%, and 20% (w/v) samples, respectively.

##### Methods for Analyzing Highly Crowded Samples

To distinguish bound from unbound probes, a tracking algorithm was employed that compared the estimated diffusion coefficient of a fluorophore within a pit to a threshold value, as described in Scott, *et al.* [2]. Briefly, the brightest pixel intensity within a pit was determined for each frame of a video. Then, the distance between the brightest pixel intensities for successive frames was calculated and compared to a threshold value: rapidly diffusing unbound probes possessed diffusion coefficients above this value, while the slowly diffusing bound probes were below. At 20% 8 kDa PEG concentrations, the diffusion of the unbound probes was measured below the threshold value for distinguishing bound and unbound probes at 50 ms exposure time. To calculate a new threshold diffusion value, samples of pUC19 were labelled with YOYO-1 intercalating dye from Thermo Fisher Scientific Canada (St. Laurent, QC) at a concentration of 1 dye per 10 base pairs. These molecules were then placed in a CLiC microscope device as per the protocol in Scott, *et al.*, and trapped in pits with 20% 8 kDa PEG, 137.5 mM NaCl, 12 mM Tris, 25 mM HEPES, 2.5 mM protocatechuic acid (PCA), 50 nM protocatechuate 3,4-dioxygenase (PCD), a pH of 8.0, and at a temperature of 37°C. Five videos with 20 frames and 50 ms exposure time were then obtained, replacing the trapped plasmid molecules between each video acquisition with ones from the surrounding solution by lifting the CLiC lens, oscillating it, and lowering it to the same height. Labelled plasmids were then tracked using the algorithm from above, and the fastest diffusion value of 0.064 $/mu\text{m}^2$/s for all the data sets

recorded. This threshold was then used when tracking the labelled oligo to distinguish a bound complex from a diffusing oligo.

#### Principal Component Analysis (PCA)

The K-NN cluster algorithm presented in Scott, *et al.* was improved upon by incorporating a technique called Principal Component Analysis (PCA). In the K-NN cluster algorithm as applied to this data, a set of predictors are chosen to determine whether a tracked probe is bound or not. These predictors are variables that are different for bound and unbound probes: the estimated diffusion coefficient, the size of the intensity spot, and location variance. For a more detailed explanation of the K-NN cluster algorithm, please refer to Scott, *et al.* [2].

PCA is a statistical procedure that uses an orthogonal transformation to change a set of possibly correlated observations into a set of values of linear, uncorrelated variables called principal components. These principal components are defined in such a way that the first principal component corresponds to the variable that accounts for the largest amount of variance, the second principal component corresponds to the variable that makes up the second largest amount of variance, and so on. In our analysis, PCA is applied to transform predictors from the data into lower dimensional representation. The predictors that account for minimal variance are then discarded in a technique called feature selection as these will have little impact on distinguishing bound from unbound probes. Since noisy or irrelevant predictors can have the same influence on the K-NN algorithm's predictions as significant predictors, they can negatively impact the accuracy of the predictions. Additionally, this prevents model overfitting, where more parameters than necessary are used to determine whether a model is valid, allowing random patterns to impact the result. Thus, feature selection improves the K-NN algorithm's predictive accuracy.

In order to determine the effects of PCA feature selection on the K-NN classifier, two K-NN classifiers were run in parallel for a single set of videos to make predictions of the number of binding events over time. One of these used PCA feature selection to remove unnecessary predictors, while the other took all predictors as input. Separately, as bound and unbound probes were distinguishable by eye, a user manually determined which pits in the system contained bound probes and which did not. Comparing the user-determined binding events with the results from the two K-NN classifier runs allowed the calculation of the true positives, where the algorithms correctly determined a bound event, true negatives, where the algorithms correctly determined no binding occurred, false positives, where the algorithms incorrectly determined binding, and false negatives, where the algorithm incorrectly determined no binding between probe and plasmid occurred. Compiling these values for each video and each K-NN algorithm run, a confusion matrix composed of the true positives, true negatives, false positives, and false negatives called confusion matrices were constructed. From the confusion matrix, the sensitivity (true positive rate) and specificity (true negative rate) were calculated, as well as the false positive and false negative rates. Sensitivity is defined as:

$$Sensitivity = \frac{TP}{P} = \frac{TP}{TP + FN} \tag{1}$$

where $TP$ are the true positives, $P$ are the total positives, and $FN$ are the false negatives. Specificity is defined as:

$$Specificity = \frac{TN}{N} = \frac{TN}{TN + FP} \tag{2}$$

where $TN$ are the true negatives, $N$ are the total negatives, and $FP$ are the false positives. High values of Sensitivity and Specificity indicate that an algorithm is suitable for determining binding events, while low values indicate either high rates of false positives, or false negatives.

The sensitivity and specificity were calculated cumulatively for an entire data set of 89 videos, as there were not enough binding events per video. Comparing the Sensitivity and Specificity, it is evident that the K-NN classifier algorithm that applied PCA Feature selection has a slightly higher sensitivity than when all predictors are used, as shown in Fig. 1.

Comparing the Specificity for the algorithm runs with and without PCA feature selection, it can be seen that the algorithm that uses PCA feature selection has a slightly higher Specificity than the algorithm that does not use it, as in Fig. 2. Since the number of true negatives vastly outnumbers the false positives for both runs, the Specificity of both algorithms is high, likely due to the large number of free probes compared to bound ones.

To quantifiably compare the ability of each algorithm to differentiate bound and unbound probes, we compared the false negative rate (FNR) and false positive rate (FPR) for each algorithm. Small values of FNR and FPR indicate better suitability at determining bound from unbound probes. FNR is defined as $FNR = 1 - Sensitivity$, while FPR can be calculated via $FPR = 1 - Specificity$. Using this methodology, the confusion matrices, the average FPR, and the average FNR were calculated for four datasets of 89 videos each ($\sim 142,400$ data points).

Comparing the two K-NN classifier algorithm runs, it is evident that when PCA feature selection is incorporated, there is a reduction in both the number of false positives and false negatives, an example of which is given for one

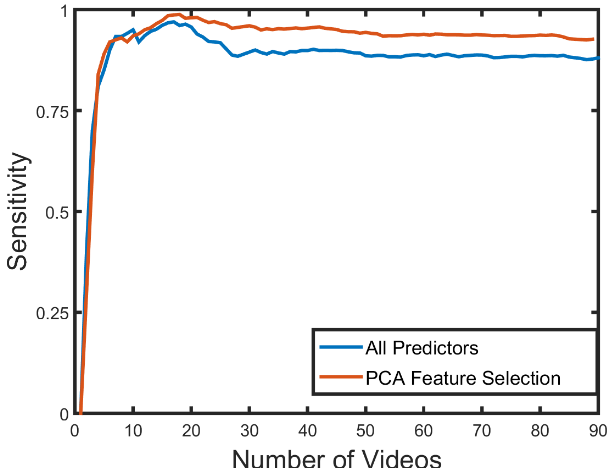


Figure 1: **Sensitivity as a function of number of videos for two parallel runs of the K-NN classifier algorithms, one with PCA feature selection, the other without.** The K-NN classifier algorithm that used all predictors to determine binding is shown in blue, while the algorithm that applied PCA feature selection is show in red. The sensitivity increases until approximately the 20th video and then plateaus there for both PCA and non-PCA algorithm runs.

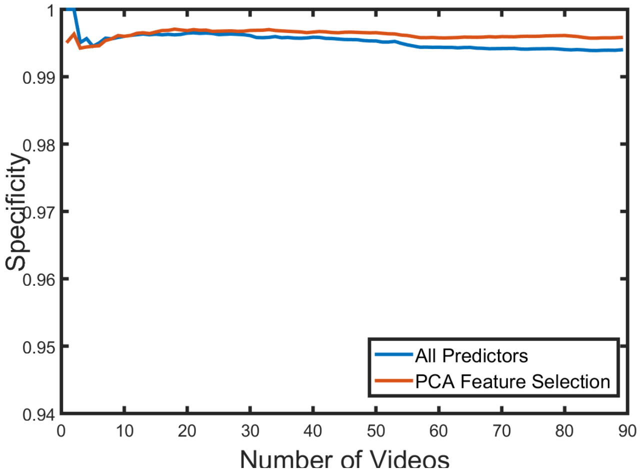


Figure 2: **Specificity as a function of number of videos for two parallel runs of the K-NN classifier algorithms, one with PCA feature selection, the other without.** The K-NN classifier algorithm that used all predictors to determine binding is shown in blue, while the algorithm that applied PCA feature selection is show in red.

data set of 89 videos in Table 1. Convincingly, both the average FNR and FPR for the four data sets are lower when PCA feature selection is incorporated, indicating an improvement on the K-NN classifier analysis. FNR, in particular, is much lower; this indicates that PCA feature selection is better at detecting binding events, leading to a lower false negative rate. Additionally, PCA feature selection also decreased the run-time of the K-NN classifier: by lowering the amount of predictors, less computation time is required to obtain results.

### List of Figures

1 **Sensitivity as a function of number of videos for two parallel runs of the K-NN classifier algorithms, one with PCA feature selection, the other without.** The K-NN classifier algorithm that used all predictors to determine binding is shown in blue, while the algorithm that applied PCA feature selection is show in red. The sensitivity increases until approximately the 20th video and then plateaus there for both PCA and non-PCA algorithm runs. . . . . . . . . . . . . . . . . 3

2 **Specificity as a function of number of videos for two parallel runs of the K-NN classifier algorithms, one with PCA feature selection, the other without.** The K-NN classifier algorithm that used all predictors to determine binding is shown in blue, while the algorithm that applied PCA feature selection is show in red. . . . . . . . . . . . . . . . . 3

| | PCA Feature Selection | All Predictors |
|---|---|---|
| False Negatives | 19 | 40 |
| True Negatives | 35394 | 35387 |
| False Positives | 59 | 66 |
| True Positives | 128 | 107 |

Table 1: **Confusion matrix for two parallel runs of the K-NN classifier algorithms, one with PCA feature selection, the other without.** This matrix was calculated for one data set composed of 89 videos.

| | PCA Feature Selection | All Predictors |
|---|---|---|
| FNR | 15.8% | 23.3% |
| FPR | 0.315% | 0.416% |

Table 2: **False negative rate (FNR) and False positive rate (FPR) for two parallel runs of the K-NN classifier algorithms, one with PCA feature selection, the other without.** These values are the average of the FNR and FPR for 4 different data sets composed of 89 videos each.

#### List of Tables

1 **Confusion matrix for two parallel runs of the K-NN classifier algorithms, one with PCA feature selection, the other without.** This matrix was calculated for one data set composed of 89 videos. . . . . . . . . . . . . . . . . . . 4

2 **False negative rate (FNR) and False positive rate (FPR) for two parallel runs of the K-NN classifier algorithms, one with PCA feature selection, the other without.** These values are the average of the FNR and FPR for 4 different data sets composed of 89 videos each. . . . . . . . . . . . . . . . . . . . . . . 4
